# Three-dimension transcriptomics maps of whole mouse embryo during organogenesis

**DOI:** 10.1101/2024.08.17.608366

**Authors:** Mengnan Cheng, Huiwen Zheng, Qi Fang, Yinqi Bai, Chao Liu, Hailin Pan, Zhewei Zhang, Huanlin Liu, Qin Lu, Chang Shi, Zehua Jing, Ning Feng, Guojun Fu, Yumei Li, Jing Feng, Tianyi Xia, Zepeng Li, Jingjing Wang, Yuanyuan Chen, Lianying Wang, Zhonghan Deng, Mei Li, Yuxiang Li, Yong Zhang, Longqi Liu, Ao Chen, Xun Xu

## Abstract

Understanding mammalian development heavily relies on classical animal models like the house mouse (*Mus musculus*). Advanced spatial transcriptomics has enabled biologists to break new ground in studies of molecular dynamics and cellular patterning during embryonic development with spatiotemporal resolution. To construct a comprehensive developmental trajectory, current three-dimensional (3D) spatial transcriptomic profiling leverages mouse embryos from gastrulation (E5.5) to organogenesis (E13.5) in continuity. However, a crucial phase for early organogenesis between E9.5 and E11.5 was deficient. To unveil the mystery of this stage, we present the 3D transcriptomics of mouse embryos at E9.5 and E11.5, with a widely applicable reconstruction workflow that bypasses sophisticated bioinformatic calculations. As our 3D atlas is generated at single-cell resolution, we demonstrate how organogenetic processes can be interpreted at different levels of granularity, from local cellular interactions to whole embryonic regionalization. We release the open-access database MOSTA3D (Mouse Organogenesis Spatiotemporal Transcriptomic Atlas in Three-dimensional) and hope a broader community will contribute to extending this framework from conception to senility in the near future.

## Introduction

Mammalian organogenesis represents a remarkable process wherein cells from all three germ layers rapidly transform into a complete embryo containing the majority of internal and external organs within a brief timeframe^1^. Much of our understanding of mammalian development relies on organs from model species, particularly the widely utilized mouse model. Mice develop swiftly from fertilization to birth in just 21 days. Following fertilization and the formation of the blastocyst by Embryonic day 4 (E4), implantation into the uterus occurs, marking the onset of gastrulation around E6.5-E7.5, establishing distinct germ layers. Subsequently, from E8-E8.5, the mouse embryo transitions from gastrulation to organogenesis, initiating lineage specification into specific organ primordia, including early neural plates and heart tubes^2^. Starting at E9.5, organ and tissue begin to form, culminating in the establishment of most organ systems. Cell numbers rapidly increase from thousands to millions during this brief window. Throughout these developmental stages, cells transition from a single germ layer lineage to diverse multicellular organ tissues through orchestrated changes in gene expression and cellular migration, ultimately forming cooperative cell populations with specific functions at precise locations. Understanding the cellular transitions and molecular mechanisms underlying organogenesis is crucial for deciphering fundamental developmental processes such as body axis establishment, organ morphogenesis, and formation of organ boundaries. Moreover, this knowledge provides insights into developmental disorders, offering potential avenues for research, intervention, and treatment strategies.

In recent years, the advancement of single-cell technologies, particularly high-throughput single-cell techniques, has revolutionized the analysis of cellular fate transitions and molecular changes during development at a global scale^3^. Large-scale single-cell atlases have meticulously defined the cell types and molecular signatures of organs at various stages of mouse embryonic development^1,4,5^. However, these approaches have traditionally lost the spatial context of the tissues. The emergence of spatial transcriptomics has addressed this gap^6^. Researchers have employed various newly developed spatial transcriptomic technologies to explore different stages of mouse embryonic development, providing a molecular cartography for more comprehensive understanding^7^. Despite these advances, most studies have concentrated on pre-organogenesis or specific time points during organogenesis. These studies are also limited with the number of curated sections, which only represent a partial view of an organ’s state. During the rapid molecular changes characterizing organogenesis, axial and gradient molecular patterns become apparent at both the embryo and organ levels. Full-scale analyses of gene expression and cell distribution at the whole-organ or even whole-embryo level can collectively shape an unbiased discovery of the molecular characteristics underlying function and structure, elucidating the critical spatiotemporal mechanisms that regulate developmental fate.

In this study, we employed Stereo-seq to generate three-dimensional spatial transcriptome maps of whole embryos during mouse organogenesis. We obtained a total of 94 slices at the E9.5 stage and 91 slices at the E11.5 stage, reconstructing comprehensive 3D atlases of two embryos with an algorithm-independent pipeline. These molecular holograms provide detailed insights into almost all organ primordia configured at this stage and recapitulate multiscale signatures from dynamic cell arrangements and migration, cell–cell communication, spatial localization and heterogeneity, to organ architecture formation during whole-embryo morphogenesis. This comprehensive data enables us to decode the complex spatiotemporal dynamics by jointly analyzing micro- and macroscopic morphogenesis signatures that drive development, serving as a valuable resource for further research in developmental biology and regenerative medicine.

## Result

### Algorithm-independent pipeline for 3D reconstruction

Current sequencing-based spatial transcriptomic experiments include the process of cryostat sectioning the sample into single-cell-thickness slices before downstream RNA capture and sequencing^8^. This cutting step, as well as subsequent mounting step, inevitably introduces tissue deformation during sample preparation. Unlike adult mouse brain samples that can be registered to the Allen Mouse Brain Common Coordinate Framework^9^, mouse embryos lose their pristine morphology even when using state-of-the-art bioinformatic algorithms for reconstruction. We developed a widely applicable pipeline to effectively address this issue by incorporating an automated Serial Block-Face Imaging (SBFI) approach (Fig1 B). SBFI is performed before the microtome blade removes each section from the surface of the specimen, regardless of whether it will be sequenced or not. The previously computational-intensive 3D reconstruction process is then streamlined by stacking the SBFI into a 3D shell and registering the spatial transcriptomics data to their corresponding block-face images (see Methods). The approach is particularly effective for slices sequenced with large depth gaps or for samples with a bending ’banana’ shape.

**Figure 1.**
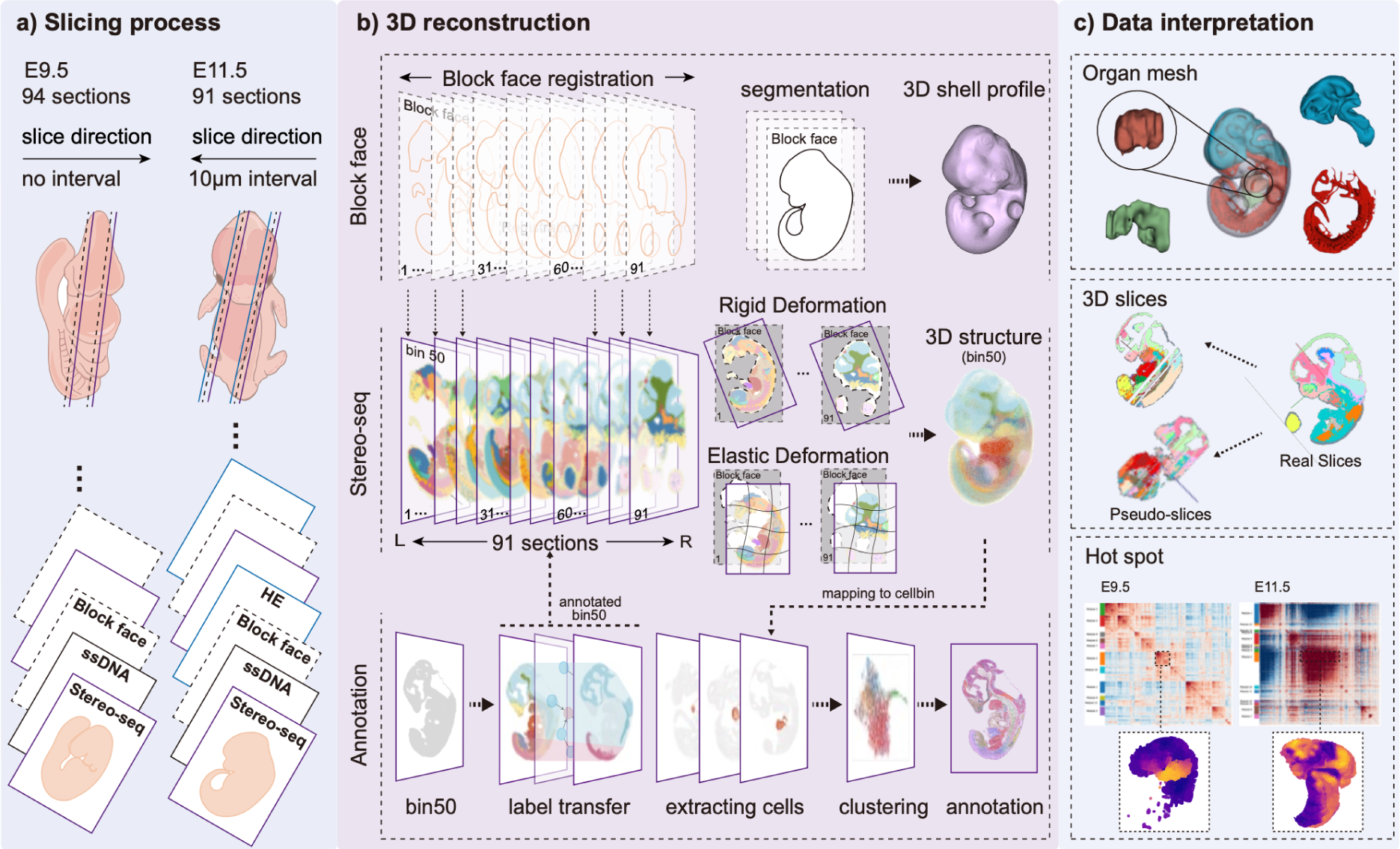
Single-cell spatial transcriptomics atlas of mouse organogenesis at E9.5 and E11.5. **a**. The detailed sectioning, imaging and sequencing processes. All sections are sequenced for the E9.5 embryo, and every other section is sequenced for the E11.5 embryo. Serial Block-Face Imaging (SBFI) is performed before the microtome blade removes each section from the surface of the specimen. ssDNA is stained for sections to be sequenced by Stereo-seq spatial transcriptomics technology, while H&E staining is applied to the neighboring sections that are not sequenced of the E11.5 embryo. **b**. The workflow of SBFI-aided reconstruction and annotation for 3D ST atlas. Each section is registered and aligned to its own block-face image to correct for nuanced sample deformation caused by the cryostat or other experimental stresses. The 3D ST data is reconstructed based on directly stacked block-face images. High-quality sections are used for spatial domain recognition at bin50 resolution, and the domain labels are transferred to neighboring sections of relatively low quality. For each 3D domain, cells are segmented and annotated with reference to the mouse embryonic single-cell atlas from Qiu et al. **c**. Downstream analyses for 3D atlas data. The mesh model defines the 3D shape of each organ. Basic statistics include organ size, cell number, and cell density. 2D pseudo-slices can be generated agnostically to the original slicing direction. Sub-domains are calculated based on spatial co-expression patterns for fine-grained organogenesis analyses.

As a result, we constructed mouse E9.5 (94 serial sections sequenced) and E11.5 (91 alternate sections sequenced) embryos through this pipeline. The consecutive tube-like neural tube (including brain and spinal cord), notochord, and gastrointestinal tract, the smooth structures of round-shaped eyes, kidneys, and limbs, as well as the 2D pseudo-slices generated agnostically to the original slicing directions, all demonstrate the accurate reconstruction effect (Fig1).

### Cell type annotation

To meticulously annotate cells, we fully leveraged gene expression profiles from each section and cross-sectional information from the reconstructed 3D mode (Fig1 B)l. We first aggregated spatial transcriptomics data into arbitrary bin50 grids (binned 50 × 50 spots into a unit grid, equivalent to a 22.5-micrometer square) for domain identification on the curated high-quality sections. The resolution of the domains was adjusted to align with known organs, and sometimes manual merging of neighboring domains was allowable to better present the structure of the organs. Accordingly, 14 organs of E9.5 and 23 organs of E11.5 were annotated, and these annotations were then transferred to the bin50s in neighboring sections of relatively lower quality using a trained graph attention network. After aggregating the same domain from all the sections, a 3D organ was created. All cells, segmented by stained ssDNA^10^, in the 3D organ were clustered using Leiden^11^ method and assigned cell type annotations to clusters by referencing the single-cell atlas of mouse organogenesis^5^ spanning E9 to E12.

Altogether, 915,901 and 7,830,602 cells were annotated into 88 and 100 cell types in E9.5 and E11.5 embryos, respectively (Fig2 A). In the E11.5 embryo, the proportion of parenchymal cells varied across different organs, ranging from 46.3% to 100%. For stromal cells, the chondrocytes (Atp1a2+), definitive erythroblasts (CD36+), early chondrocytes and fibroblasts were the most widely detected across the organs, ranging from 0.8% in dorsal root ganglion for Definitive erythroblasts (CD36+), to 22.9% in sclerotome for early chondrocytes. We showcase three representative organs *in situ*, including the spinal cord, heart, and brain, illustrating how cells are orchestrated into sophisticated structures in the molecular hologram (Fig2 B-E).

**Figure 2.**
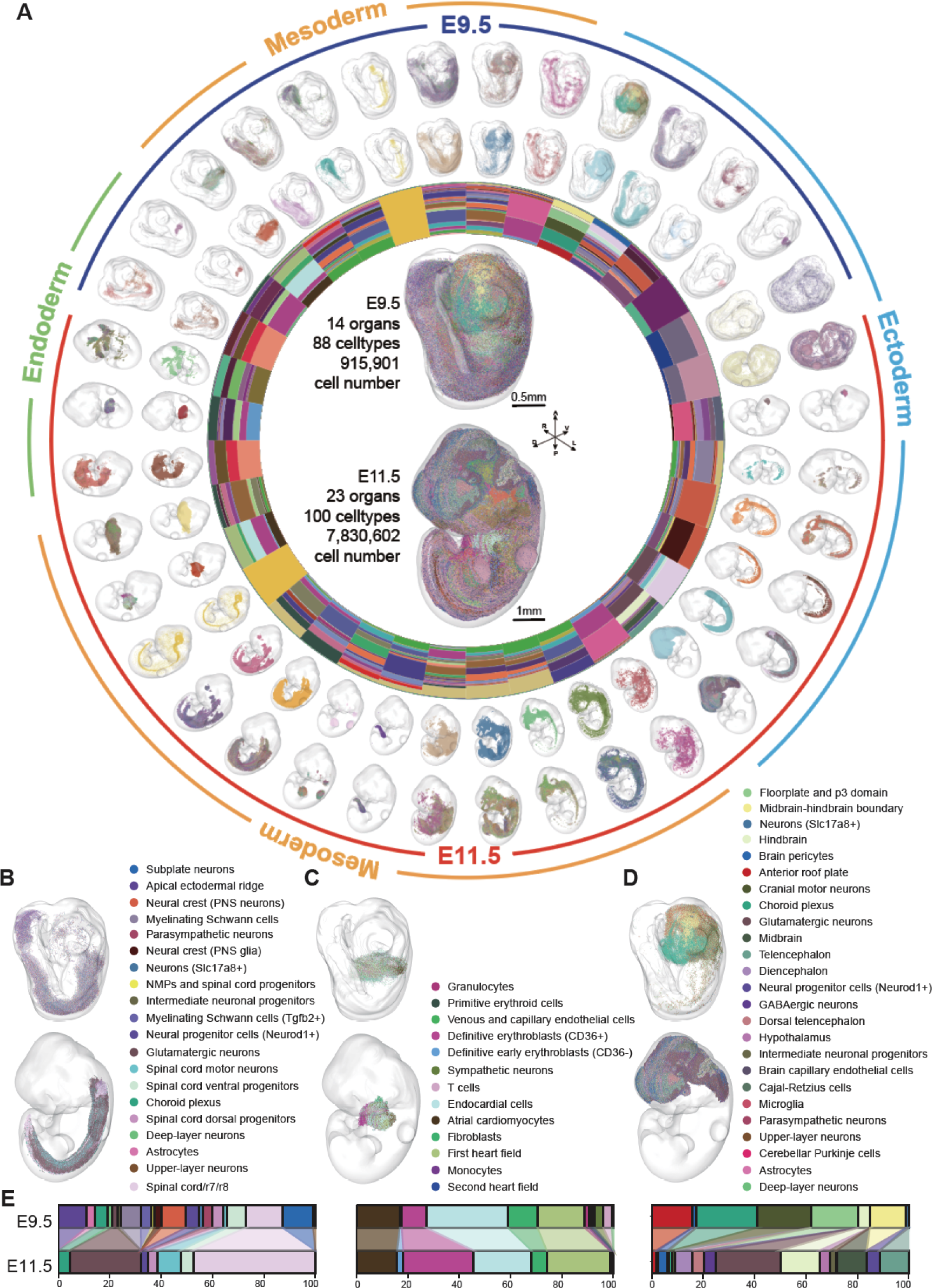
Overview of 3D cell cartography for E9.5 and E11.5 mouse embryos. **a**. Concentric circles chart showing the fine-grained cell type compositions (**i**), the *in situ* distribution of major cell types (**ii**), and the *in situ* distributions of organs (**iii**) for E9.5 and E11.5 embryos. **b-d**. Visualization of cell type distributions for representative organs, including spinal cord (**b**), heart (**c**), and brain (**d**). **e**. The Diagram shows the comparison of cell ratio of spinal cord (left), heart (middle) and brain (right y axis) between E9.5 and E 11.5 embryos.

The dynamic variations between these two crucial phases of organogenesis is also of interest (Fig3 A). The nearly tenfold increase in total cell number is mainly contributed by the proliferation of glutamatergic neurons and sclerotome cells. Besides the emergence of the otic vesicle and the neural crest, the organs that expanded most in size are liver, gut and brain. On the other hand, the overall cell density remains similar or slightly decreases across most of the organs, demonstrating a negative correlation between these two metrics during development (Fig3 B). The spatial rearrangement, differentiation, and migration of cells are the major reasons for this phenomenon. Thus, we next analyze the potential molecular driving factors for the corresponding morphogenesis.

**Figure 3.**
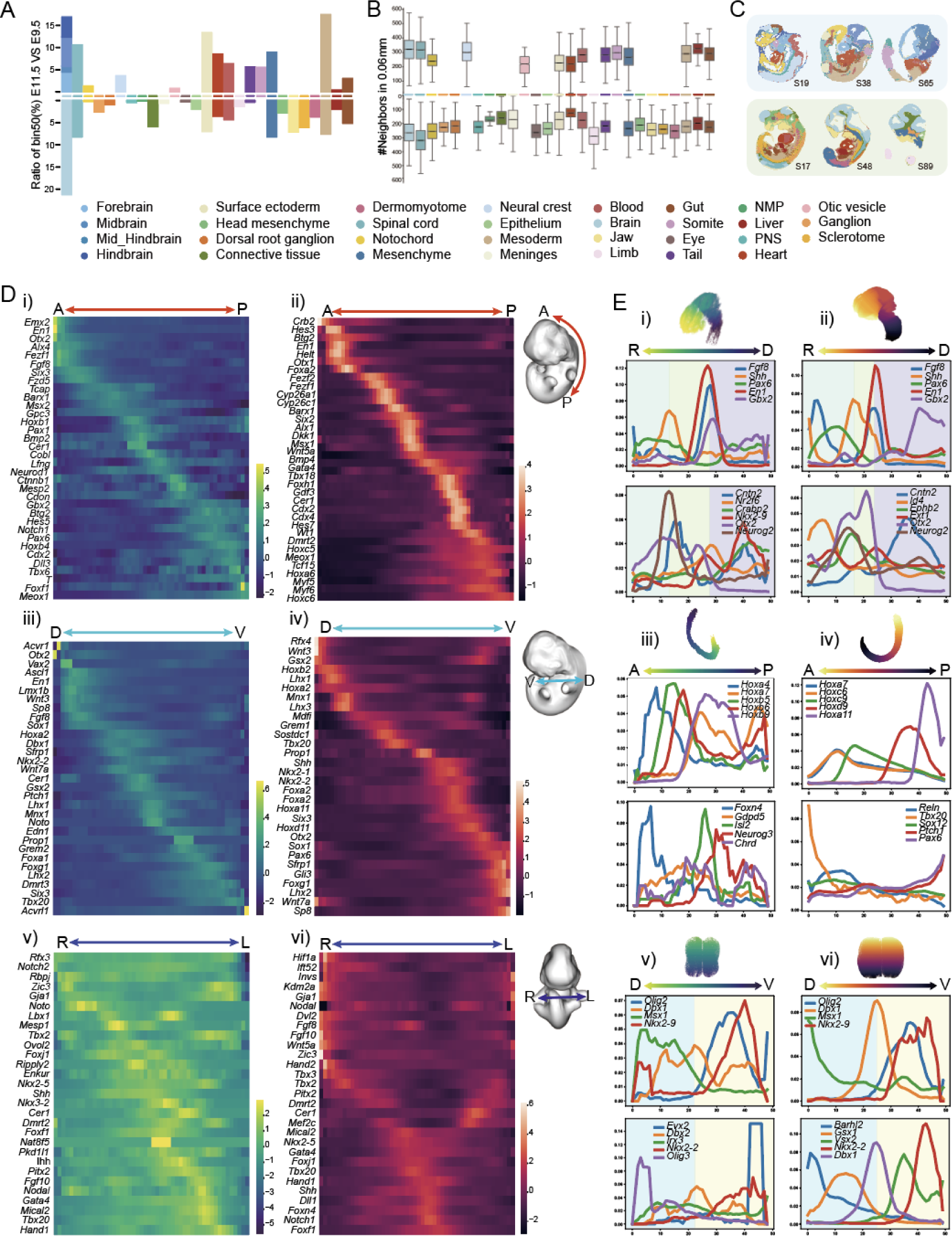
Transcriptional differences on the developmental axes between E9.5 and E11.5 embryos. **a**. Bi-directional bar chart of the changes in organ volume. Organs are sorted by the fold change in expansion size between E9.5 and E11.5. **b**. Boxplot of sorted cell density changes among the organs. **c**. Visualization of representative sections showing the organ distributions on the left, middle, and right sagittal planes. **d**. The heatmap of genes that are highly variable along the anterior-posterior (**i** and **ii**), dorsal-ventral (**iii** and **iv**), and right-left (**v** and **vi**) axes of the entire E9.5 and E11.5 embryos, respectively. **e**. Density plot of the distribution along the rostral-caudal axis (**i** and **ii**) of E9.5 and E11.5 brain, the anterior-posterior (**iii** and **iv**) and dorsal-ventral (**v** and **vi**) axes of E9.5 and E11.5 spinal cord.

### Spatial and temporal regionalization

Until now, how an organ or a system develops diverse interconnected cell types from a few stem cells has remained a challenging question. The spatial and temporal patterning mechanism, also known as embryonic regionalization process, provides a valuable model through which diversity can be generated^12^. By implementing a 3D digitization method in our Spateo package, we are able to explore the distinct sets of morphogens, transcription factors, and other molecular signatures that define the distinct cell distributions along orthogonal axes in both spatial domains and temporal windows for the entire embryo.

We first explore and validate a cascade of documented axis-patterned genes at E9.5, and then examine their expression profiles at E11.5 to determine how the graded distribution is maintained or shifted during this developmental phase (Fig3 D). Conversely, the E11.5 patterned genes are traced back to their distributions at E9.5 for the same purpose. Subsequently, we attempt to identify candidate genes that strictly follow the axis patterns of known genes, considering them as potential novel biomarkers for pattern formation during organogenesis.

On the anterior-posterior axis, 240 genes (corresponding GO term: 0009952) are curated for this analysis (Fig3 D). At the stage of E9.5, the representative regionalization genes are expressed from the forebrain (e.g., *Emx2*, *Otx2*) to the caudal region (e.g., T), most of which are classical transcription factors (TFs). Strikingly, more than half of these TFs no longer govern the same region in the regionalization process at E11.5, indicating the cell differentiation is sustained locally and transiently while TFs are versatile during development in this period. For example, *Emx2* (the first gene along the A-P axis at E9.5), in cooperation with *Otx2* (the third one), is known to generate the boundary between the roof and archipallium in the developing brain. While it is recently reported to be transiently expressed at the base of the limb buds, as well as in mesenchymal cells of the Wolffian ridge of E11.5 embryos^13^. This phenomenon also recur on the dorsal-ventral and medio-lateral (a.k.a. left-right) axes.

In addition to the overall embryonic axial gene expression, we also examined axial gene expression in the brain and spinal cord (Fig3 E). Brain morphogenesis at this time is coordinated by intrinsic signals from neural progenitors, through which we distinguish the forebrain, midbrain, and hindbrain along the rostral-caudal axis of the brain^14^. These signals include *Fgf8*, which is emitted by the anterior neural ridge (ANR), and is also elevated at the junction of the midbrain and hindbrain by the isthmic organizer (Iso). Also elevated here are *En1* and *Gbx2*, and SHH signals emitted by the zona limitans intrathalamica (ZLI) at the junction of the telencephalon and diencephalon.

We further distinguished neuron-related signals with the same trend under the corresponding modules. *Cntn2* is a cell adhesion molecule of the contactin family of neuronal recognition molecules, which is involved in neuronal migration, axon guidance, and the organization of myelin subdomains^15^. The signal was more significant in the hindbrain at E11.5 than in the previous period. Next, we observed the distribution of the Hox gene family on the A-P axis of the spinal cord. It was surprising that *Hoxa7* and *Hoxc6* had consistent fluctuations^16^. On the D-V axis of the spinal cord, the dorsal and ventral signals were divided with *Dbx1* as the boundary. The corresponding neuron-related signals were still relatively unstable at E9.5, and had shown regional significance at E11.5.

### The increasingly complex spatial expression patterns of the nervous system

From E9.5 to E11.5, the mouse nervous system undergoes significant developmental changes, including the subdivision of brain vesicles, the formation of cortical neurons, and the development of the eyes^2^. At E9.5, the nervous system is in an early stage of development, characterized by broad and general gene expression patterns. Our data reveal widespread signals related to cell division (E9.5 brain module 7, Fig4 A) throughout E9.5 brains (Fig4 B left), with high gene set scores for cell division (GO:0051301) also present in axon growth domains (E9.5 brain module 5). We did not identify significantly differentially expressed cell cycle-related genes between these two modules. While axon growth domains have more genes involved in the regulation of neural progenitor proliferation, including *Nr2f1, Sfrp1, Sema5b*, and *Hes5*^17–20^. However, by E11.5, the distinction between cell division domains (E11.5 brain module 2) and axon growth domains (E11.5 brain module 1) becomes more pronounced, as indicated by spatial co-expression patterns and gene set scores (Fig4 C-D left). The cell division regions at E11.5 are enriched with several cell cycle-related genes, including *Top2a, Ccnd1, Cenpa*, and *Cdca8*^21–24^. The shift from broad, general signals to more localized and specialized patterns indicates the rapid maturation of the neural system, which is consistent with the decreased expression fraction of specific genes in cells of these two modules (Fig4C-D left, dot plots).

**Figure 4.**
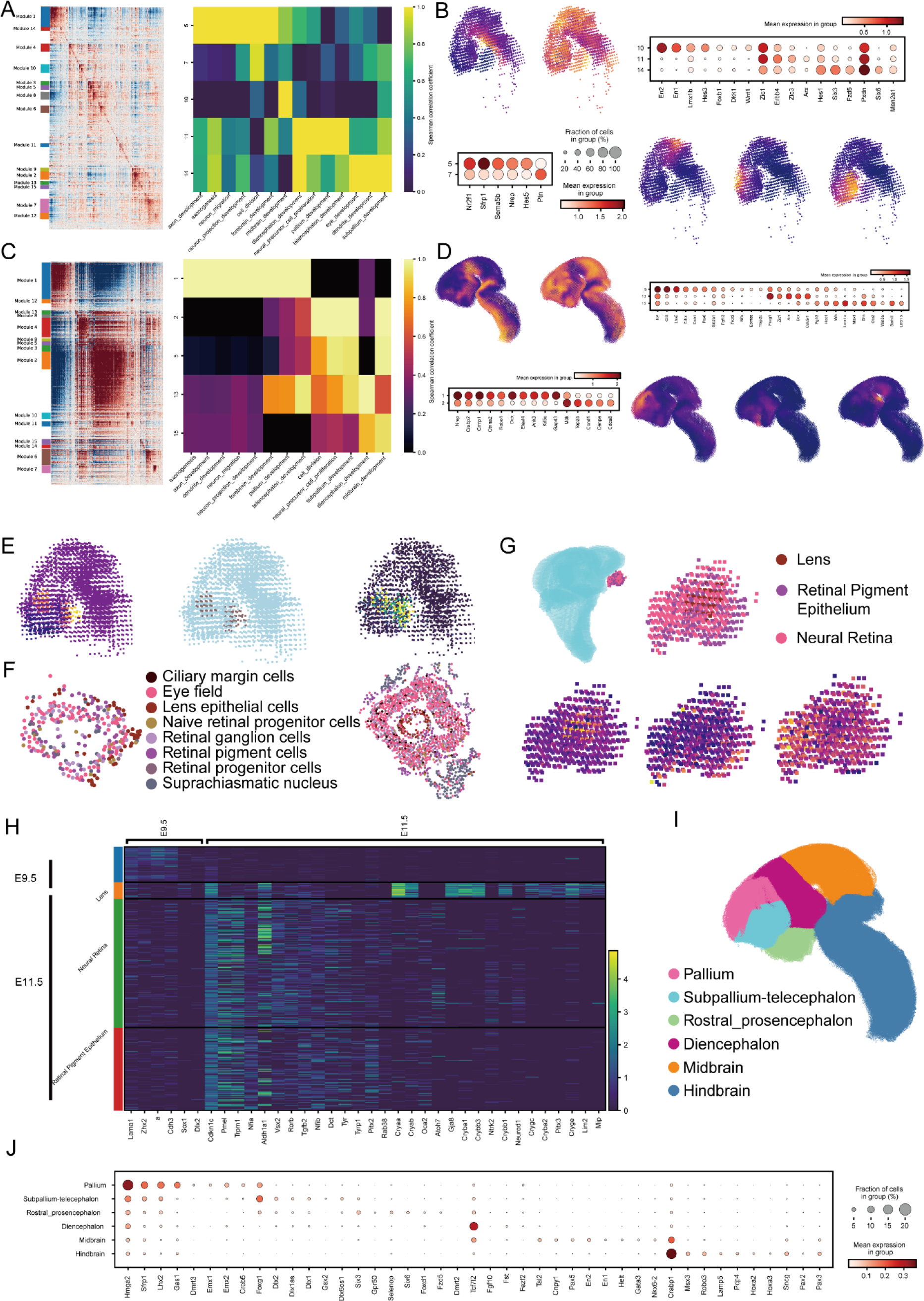
Developmental changes in the mouse brain lead to increasing spatial complexity. **a**. Hotspot result (**Left**) of the identified spatial co-expression modules (a.k.a. domains) at E9.5. The enrichment analysis (**Right**) showing the main GO terms of the selected domains. **b**. In situ visualization of each selected domain. Bubble plot showing the expression of marker genes in axonogenesis and cell division related domains (**Left**), as well as midbrain, pallium, and subpallium related domains (**Right**). **c**. Hotspot result (**Left**) of the identified spatial domains at E11.5. The enrichment analysis (**Right**) showing the main GO terms of the domains that corresponds to E9.5. **d**. Spatial locations and marker genes of corresponding domains at E11.5. **e**. Fine-grained hotspot result showing (**Left**) the early development of eye at E9.5, verified by cell type annotation (**Middle**) and marker gene expression (**Right**). **f**. Distribution of cell types of one single eye at E9.5 (**Left**) and E11.5 (**Middle**). **g**. The lens, retina, and retinal pigment structure of an eye. **h**. Differential expressed genes between E9.5 and E11.5. **i**. schematic of brain areas at E11.5. **j**. Bubble plot showing the expression of marker genes in different brain areas.

With the progression of development, in addition to co-expressed genes related to neural progenitor differentiation and proliferation (for example *Nrep, Crabp2* and *Elavl4*)^25,26^, we also identify various co-expressed genes that regulate axon guidance and neuronal migration which is important to the formation of neural circuits, such as *Crmp1, Ctnna2, Robo1* and *Dcx*^27–30^.

From E9.5 to E11.5, axon growth signals expand from the ventral regions of the mid-hindbrain and spinal cord to broader areas of the brain, which is also indicating increased neuron migration and the establishment of specific cortical architecture.Analysis of spatial co-expression modules associated with brain vesicles reveals the increasing complexity of brain development.

Although only three primary brain vesicles (forebrain, midbrain, and hindbrain) are formed by E9.5, our spatial transcriptomic data (Fig4 A-B) identifies distinct co-expression patterns between the pallium and sub-pallium within the telencephalon (E9.5 modules 11 and 14), showing ongoing development process. At E9.5, signals for maintaining neuronal progenitors, such as *Zic1* and *Zic3*^31^, are more pronounced in the pallium. Additionally, we observe the co-expression of *ErbB4* (a direct target of ARX) with *Arx*, which is involved in the development of cortical interneurons migrating from the ganglionic eminence to the cortex^32^. While previous studies suggest the formation of the ganglionic eminence begins around E10-12^2,33^, our spatial co-expression data indicate earlier signals of interneuron migration within the cortical cortex (ependymal layer).

By E11.5, the nervous system exhibits increased complexity and more subdivisions, with the cerebral neocortex beginning to differentiate into layers. We then manually segment distinct brain regions based on co-expression modules and anatomical structures, including the pallium, subpallium-telencephalon, rostral prosencephalon, diencephalon, midbrain and hindbrain (Fig4 I). With the differentially expressed genes (DEGs), we identify many key regulatory genes in each brain region, such as *Tcf7l2*, which maintains thalamic neurons in the telencephalon^34^; *En1*, which promotes midbrain development; *En2*, which promotes cerebellar formation^35^; Hox genes, which define the hindbrain patterning^36^; and Emx genes, which are involved in cortical subdivision^37^ (Fig4 J). In pallium (E11.5 module 5), we observe high gene set scores of neural precursor cell proliferation (GO:0061351) (Fig4 C right). At E11.5, we show higher expression of *Pax6* and *Eomes* in pallium(Fig4 D right), while sequentially expressing *Pax6*, *Tbr2* (also known as *Eomes*), and *Tbr1* determine the fate of two precursor cells in cerebral cortex: intermediate progenitor cells and radial glia (the progenitors of projection neurons)^38^. *Pax6*, and *Eomes* are also enriched in intermediate progenitor cells of pyramidal neurons, which are derived from radial glial cells that converted from neuroepithelial cells in the ependymal layer at E9.5^39^. Additionally, various pyramidal neurons, differentiated from radial glial progenitors, are crucial for forming the neuronal network in the cerebral cortex. This process is regulated by the expression of *Lhx2* and *Fezf2*, both of which are enriched within the pallium in our spatial data^40^. Our spatial transcriptomic data illustrate the gene co-expression patterns of the cerebral cortex formation, revealing the active processes of proliferation, differentiation, and migration of various neural progenitor cells in the 3D space.

During E9.5 to E11.5, the optic vesicles undergo further differentiation, leading to the lens formation and the retinal pigmentation developments. By annotating spatial domains, we identify eyes at E11.5 but not at E9.5 (Fig2 A). However, we observe strong signals of eye development (GO:0001654) in the subpallium region at E9.5(Fig4 A right, E9.5 brain module 14). To further investigate, we conducted a second round of spatial co-expression analysis using Hotspot, which revealed a new module enriched for eye development (Fig4 E left). Then this new module is segmented as the eyes at E9.5 (Fig4 E middle). To validate these results, we examine the spatial distribution of *Rax*, a gene specifically expressed in the optic vesicles, the optic stalk, and the ventral diencephalon at E9.5 (Fig4 E right)^41^.

With cell type annotations, we illustrate single sagittal sections at E9.5 (section 10_2) and E11.5 (section 09) as shown on the left side of Fig4 F. At E9.5 the structure of the optic vesicles is clearly visible, but the lens epithelial cells are still surrounding the eye field and retinal cells, which have not yet invaginated to form the lens. In contrast, by E11.5, the eye exhibits a mature lens and a more organized retina (Fig4 F right). The distribution of cells also suggests the transition from a simpler physiological structure to a more regular and complex one. In the whole embryo, we observe a distinct separation of the eye from the forebrain at E11.5 (Fig4 G), with the intervening space filled by head mesenchyme (not shown in main figure). When identifying spatial co-expression modules within the E11.5 eye, we recognize a distinct lens module, localized to the center of the eye (Fig4 G). In addition, we annotate a module with high signals for retinal pigment epithelium development (GO:0003406) near the body surface, and another module with high signals for neural retina development (GO:0003407) near the forebrain (Fig4 G lower panel)^42^.

With the spatial transcriptomic data, we identify differentially expressed genes between E9.5 and E11.5 (Fig4 H). At E9.5, the DEGs primarily represent general eye development signals. For example, *Lama1* is expressed in the inner limiting membrane within the eyes^43^; *Zhx2*, a transcriptional regulator of *Pax6* plays a role in optic vesicles development^44^; *Cdh3* is expressed in optic cup^45^; and *Dlx2* is found in retinal ganglion cells^46^. During this stage, the lens placode is in formation with the expression of *Six1*^47^. After the lens vesicle closes at E11.5, *Six1* becomes restricted to the lens, where crystallin proteins essential for lens function are all enriched, as revealed by our spatial data (Fig4 H). Interestingly, we also observe higher expression of the nonagouti gene (*a*) at E9.5, which activates melanocortin receptor binding^48^. The role of nonagouti in subsequent melanin production remains unclear, but we note higher expression of *Tyr* and *Oca2* at E11.5, which are associated with retinal pigmentation development^49^. Except for lens and retinal pigmentation Epithelium, the neural retina identified in our spatial co-expression data at E11.5 shows elevated signals of neurons in the eyes. For instance, *Vsx2* (also known as *Chx10*) is a key transcription factor for retinal progenitor proliferation^50^; *Atoh7* is essential for retinal ganglion cell development^51^; *Ntrk2*, a receptor for brain-derived neurotrophic factor (BDNF) is involved in retinal ganglion cell survival^52^; and *Neurod1* regulates the differentiation of photoreceptors^53^. These findings highlight the more advanced and organized developmental patterns of the eyes at E11.5.

### The difference between left and right lung

At E9.5, the gut tube entered the morphogenesis stage, where organ buds of endodermal origin began to emerge. Next, we explored functional enrichment within specific regions of the gut tube using Hotspot, which integrates spatial location information with gene expression data.These include the lung domain, characterized by signals for lung and respiratory system development; the pancreas domain, marked by signals for pancreatic development; the gut domain; the pharyngeal region located at the upper end of the gut tube; and the kidney tissue located at the lower end. The kidney domain expressed the genes Hoxd10 and Hoxd11, the lung domain expressed the lung bud marker gene Nkx2-1, and the pancreas domain expressed the characteristic genes Pdx1 and Ptf1a. (Fig5 A-B)

**Figure 5.**
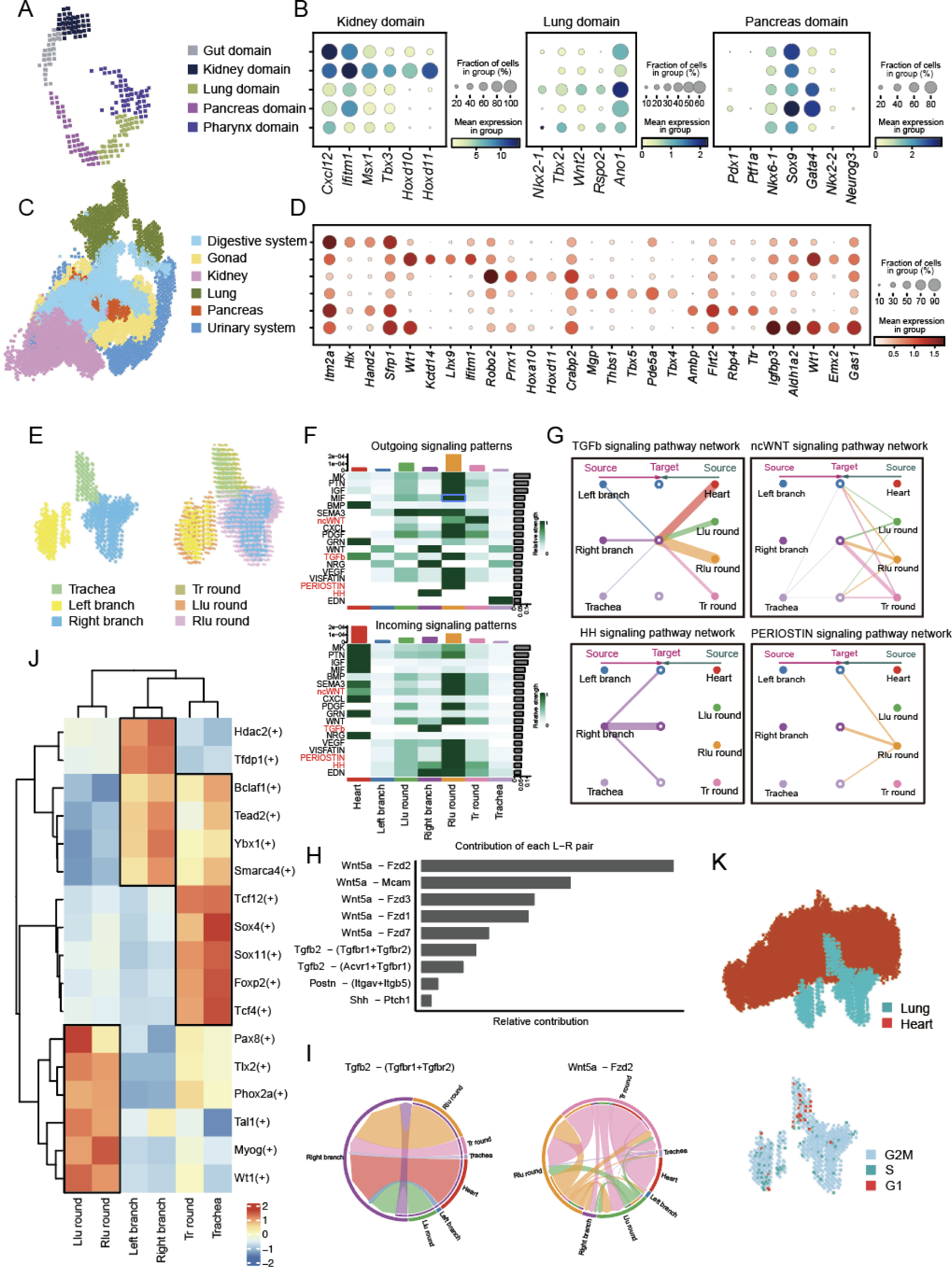
Left-right pattern of gene expression in mouse lung. **a.** E9.5 gut gene co-expression module division and annotation. **b.** Bubble plot of marker genes corresponding to the modules in Figure **a. c.** E11.5 gut and mesoderm gene co-expression module division and annotation. d. Bubble plot of highly variable genes corresponding to the modules in Figure **c. e.**The structure of the lung and the tissues surrounding them. **f.** Distribution patterns and strengths of input and output signals in seven tissues. **g.** The network of different signaling pathways acting on lung tissue itself through autocrine or paracrine. h. The contribution of ligand-receptor pairs to the signaling pathway in Figure **g. i.** Circle diagram of the interaction strength of ligand-receptor pairs. **j.** Standard deviation of the active regulons. **k.** Spatial distribution of lungs and heart (above), cell cycle phase distribution in the lung (below).

For the E11.5 samples, we extracted gut tube and mesodermal tissues for Hotspot analysis to better identify the source of the endoderm and the source organ of the mesoderm. We identified a total of six organs, corresponding to the digestive system, gonads, kidneys, lungs, pancreas, and urinary system. The spatial distribution of these tissues is illustrated in Figure 5(C). By this stage, the differential genes for these tissues were clearly expressed, unlike at E9.5, where their expression was more scattered throughout the gut tube. The kidney domain at E11.5 remained consistent with that at E9.5, marked by genes such as Ttr and Hoxd10. Additionally, Crabp2 was highly expressed in the E9.5 pancreas. (Fig5 C-D)

The morphology of the lung tissue observed aligns well with previously reported findings. The trachea bifurcates into the left and right lungs, with the left lung being less clearly defined and likely featuring a terminal bud. In the right lung, distinct lobes are visible: the cranial lobe extending forward, the medial lobe extending laterally, the accessory lobe located posteriorly, and the caudal lobe below. Notably, no buds are visible on the caudal lobe at this stage.

While the differentiation between the left and right lung branches occurs around E9.5 or E10.5, questions remain regarding whether there are differences in the subsequent development of the left and right lungs after the formation of the lobe branches^54^. Mesenchymal signals play a crucial role in guiding epithelial cell budding, branch morphogenesis, and the structural and functional maturation of the lung^55^. The impact of surrounding mesenchymal signals on the trachea, right lung branch, and left lung branch is explored in this context. Additionally, given the heart’s significant role in lung development and its inherent left-right morphological asymmetry, the heart is also considered in this analysis. The mesenchymal regions surrounding these three lung components are distinguished according to their spatial positions, left lung round (Llu round), right lung round (Rlu round), trachea round (tr round) respectively. (Fig5 E)

We used CellChat to analyze the coordination of input and output signaling pathways in seven tissue regions. The output signaling patterns of the Rlu and Llu regions are generally consistent, though differences in signal strength were observed. Notably, PERIOSTIN signaling in the Rlu region differs significantly from that in the Llu region. Among the signaling pathways analyzed, both ncWNT and TGFb signaling are reported to play crucial roles in lung development. The HH signaling pathway, which is essential for lung branch formation, shows significant activity in the right lung branch pattern. In terms of incoming signaling patterns, TGFb signals are more prominent in the right lung branch. (Fig5 F)

The network of the four signaling pathways highlighted in the figure will be presented next. The right side represents paracrine signals, while the left side represents autocrine signals. We focus on the pathways from Llu round to the left branch and Rlu round to the right branch. Although there are similar connections from Tr round to the right branch, this suggests that ligand expression related to these pathways also occurs in Tr round.In the TGFb signaling pathway, no direct link from Llu round to the left branch is shown. However, the expression of ligand and receptor genes (e.g., Tgfb2, Tgfbr1) in this pathway is not low, but rather relatively high in Rlu round and the right branch (see supplement). The ncWNT signaling pathway shows similar intensity in both the left and right lungs. The HH signaling pathway is primarily mediated by Shh-Ptch1. The network reveals that the right branch engages in autocrine signaling on itself, while no corresponding signal is observed in the left branch. Gene expression analysis supports this, as Ptch1 expression in Shh is indeed high in the right branch. Additionally, the network for the PERIOSTIN signaling pathway shows links from Rlu round to the right branch. Periostin is known to play a significant role in airway development and alveolar epithelial repair^56^. The above results show that there are quantitative differences in signal reception and secretion between the left branch and the right branch. (Fig5 F-I)

Additionally, we used Scenic software to identify differences in regulatory networks in six regions other than the heart. The Tr round and Trachea exhibited similar patterns. The transcription factor modules in the left and right branches are identical, but the scores are higher in the right branch, particularly for factors like Hdac. Furthermore, Pax8 scores higher in the Llu round compared to the Rlu round. Overall, there are notable differences in signaling pathways, gene expression, and transcription factor regulation modules between the left and right lung branches.

During this stage, the lung is undergoing lobe branching and transitioning into domain branching. Cell cycle analysis shows that a relatively higher proportion of bins in the trachea are in the G1 phase, with fewer bins in the lung lobes, where most bins are in the more active G2M phase. Additionally, there are more bins in the S phase in the left lung than in the right lung. We hypothesize that the left and right lungs are at different stages of cell cycle progression, which may contribute to the observed differences in signaling.

## Discussion

With the advances in sequencing-based spatial transcriptomics (sST) to cellular resolution, we profiled the 3D transcriptome of 0.9 million cells from E9.5 and 7.8 million cells from E11.5 mouse embryos. The total number of cells approaches the 12-million single-cell atlas that was curated from 83 staged embryos. Combined with this whole-embryo scRNA-seq atlas from Jay’s lab, we constructed a molecular hologram to present the crucial phase of mouse organogenesis. Although the RNA capture efficiency in sST is lower than in scRNA-seq data, the preserved spatial patterning and increased cell number significantly improve interpretability and reduce bias in our dataset. For example, unlike the inference of gene expression along the trajectory of similarity-based cell embedding, the gradient and regionalized patterning of transcription factors and other molecular signatures are more straightforwardly exhibited in situ. Most of our observations are manually verified by ISH results on the eMouseAtlas website to enhance the credibility of the evidence for the novel findings that are deficient in staining.

Our goal is not only to provide data resources but also to construct a general experimental 3D transcriptomic framework for the community. During cryostat sectioning, a fixed camera is mounted directly in front of the embedded sample block to photograph each section before cutting. This SBFI strategy is widely used for image analysis of scanning electron microscopy. Now, it can be seamlessly applied to sST studies, as we demonstrated on two whole-embryo mouse models. Currently, due to the high cost of sST experiments, most 3D projects are limited by large depth gaps in adults or late-stage embryos, which makes 3D reconstruction for sST data more challenging. Additionally, occasional low-quality sections can undermine the entire experiment because most reconstruction processes rely on sectional stacking. In light of image engineering, our pipeline bypasses all sophisticated bioinformatic algorithms for accurate reconstruction. For samples without SBFI or those unable to establish SBFI, we recommend our Spateo package to perform high-quality 3D reconstruction. Spateo is a specialized analytical 3D transcriptomic framework published as our companion paper. We also implemented its 3D digitization algorithm to discover novel genes that follow the axis patterning in this research.

Left-right asymmetry is an essential feature in bilateral animals^57^. The mechanism underlying the left-right asymmetrical organ morphogenesis is a central question in developmental biology. In this study, we took a first glance at left-right asymmetrical gene expression at the whole embryo scale and then linked the potential kinetics to asymmetrical organ morphogenesis, such as lung development. Though it is an initial attempt to bridge micro-scale gene alterations to macro-scale morphogenesis, we think the concept of the computational framework deserves generalization.

From gastrulation (E5.5 to E7.5^58^, and E8.5^59^) to organogenesis (E9^59^ and E13.5^60^), mouse embryonic 3D transcriptomics are emerged to advance the foundation for understanding mammalian embryonic development. Our research at E9.5 and E11.5 successfully fills the gap in early stages of organogenesis. Although these datasets are from different individuals or even different platforms, they are bridging a ground-truth molecular hologram of the prenatal development model for the community. Rome wasn’t built in a day; more data and analytical tools are anticipated in this field, and we hope a broader community will contribute to extending this framework from conception to senility in the near future.

## DATA AVAILABILITY

Spatial embryo transcriptomic data has been deposited into CNGB Sequence Archive (CNSA)^61^ of China National GeneBank DataBase (CNGBdb)^62^ with accession number CNP0005981, which has not yet been made public. The processed data is uploaded to the MOSTA-3D website (https://db.cngb.org/stomics/mosta/3d/).

## METHOD DETAILS

### Tissue imaging and processing

#### Blockface processing

We use Blockface image acquisition devices for tissue imaging, E11.5 uses 0.65× magnification lens, and E9.5 uses 2× fixed focus lens. Based on the SDK camera software system provided by Hikvision camera, the image status is adjusted, and the image clarity is ensured by adjusting the interface brightness, exposure value and gain parameters. Starting from the first slice, the image number is recorded as S1, and image acquisition is performed after each slice until all tissues are cut.

#### Tissue processing

When the whole tissue is exposed, the tissue slice is pasted to the chip, and blockface photography and pasting are performed alternately. Tissues were processed using conventional Stereo-seq workflows.Adhere the tissue sections to the surface of the Stereo-seq chip and incubate at 37°C for 3-5 minutes. Then, fix the sections in methanol and incubate at -20°C for 40 minutes before proceeding with Stereo-seq library preparation. If instructed, stain the same sections with a nucleic acid dye (Thermo Fisher, Q10212) and image them using a Ti-7 Nikon Eclipse microscope, followed by in situ capture in the FITC channel.

### Stereo-seq library preparation and sequencing

Tissue sections were initially washed with 0.1× SSC buffer (Thermo, AM9770) containing 0.05 U/ml RNase inhibitor (NEB, M0314L) to remove photographic staining. The sections were then permeabilized using 0.1% pepsin (Sigma, P7000) in 0.01 M HCl buffer at 37°C for 6 minutes. After this, the sections were washed again with 0.1× SSC buffer supplemented with 0.05 U/ml RNase inhibitor.

Reverse transcription was carried out at 42°C for 3 hours with Superscript II (Invitrogen, 18064-014). The reaction mix included 10 U/ml reverse transcriptase, 1 mM dNTPs, 1 M betaine solution PCR reagent, 2.5 mM Stereo-seq-TSO, 1× First-Strand buffer, 7.5 mM MgCl2, 5 mM DTT, and 2 U/ml RNase inhibitor. After reverse transcription, the tissue sections were washed twice with 0.1× SSC buffer and digested with tissue removal buffer (10 mM Tris-HCl, 25 mM EDTA, 100 mM NaCl, 0.5% SDS) at 37°C for 30 minutes.

The chip containing cDNA was then treated with exonuclease I (NEB, M0293L) at 37°C for 1 hour to cleave the cDNA from the chip. The cDNA was subsequently collected and washed once with 0.1× SSC buffer.

The collected cDNA was amplified using KAPA HiFi Hotstart Ready Mix (Roche, KK2602) and 0.8 mM cDNA-PCR primers. The PCR program consisted of an initial incubation at 95°C for 5 minutes, followed by 18 cycles of 98°C for 20 seconds, 58°C for 20 seconds, and 72°C for 3 minutes. A final extension step was performed at 72°C for 5 minutes.

For library construction, 20 ng of cDNA was used. Fragmentation was performed using in-house Tn5 transposase at 55°C for 10 minutes. The reaction was then terminated by adding 0.02% SDS buffer, followed by gentle mixing at 37°C for 5 minutes. The fragmented cDNA was combined with the following components for amplification: 25 μL of the fragmented product, 0.3 μM Stereo-Library-F primer, 0.3 μM Stereo-Library-R primer, and 1× KAPA HiFi HotStart Ready Mix, with the final volume adjusted to 100 μL by adding nuclease-free H2O. The amplification process involved an initial denaturation at 95°C for 5 minutes, followed by 13 cycles of 98°C for 20 seconds, 58°C for 20 seconds, and 72°C for 30 seconds, with a final extension at 72°C for 5 minutes. The amplified PCR products were purified using Ampure XP Beads (Vazyme; 0.6× and 0.2×) to generate DNBs, which were then sequenced as paired-end 100 bp reads on an MGI DNBSEQ-Tx sequencer.

### Preprocessing of Stereo-seq Sequenced Reads

Stereo-seq raw data are paired-end reads, with read 1 and read 2. In detail, read 1 consists of the coordinate identifier (CID) and molecular identifier (MID, equivalent to UMI), while read 2 contains the captured cDNA sequences.

Raw reads are processed using the Stereo-seq Analysis Workflow (SAW)^63^ to generate gene count matrices in *.gef format. Briefly, cDNA sequences are aligned to the reference genome (mm10) and filtered with MAPQ ≥ 10. Coordinates of sequenced reads are obtained by mapping CIDs to the chip. The gene expression matrix is generated using exonic reads with CID information. The resulting gene count matrix contains gene ID, x and y coordinates on the chip, MID count, exonic count (indicating completely spliced RNA molecules), and intronic count (indicating incompletely spliced or unspliced RNAs). The *.gef files are subsequently converted to *.gem format using geftools to facilitate downstream analyses.

### Cell Segmentation

To identify the location of cells in spatial transcriptomic data, we first register ssDNA images to gene count matrices which are visualized as images. An in-house Python script extracted x and y coordinates from gene count matrices along with corresponding MID counts, constructed sparse matrices representing the expression data, and saved them as .tiff images. Each .tiff image represents the spatial distribution of gene expression levels in a given section. Corresponding gene expression and ssDNA images of the same section are rigidly transformed (including isotropic scaling, translation, and rotation) manually with track lines (3-pixel wide, indicating 1.5 μm) on the two images to align them. The pipeline allows for a maximum registration error of 5 pixels.

Cell segmentation is performed using spatial positions of stained nuclei from ssDNA images using Cellbin^64^. Sequential integer labels are assigned to individual cell masks and 0 to the background. A .tif formatted cell mask image is generated, and cell labels are added as one new column to the .gem files.

### Tissue Region Segmentation

Tissue region is identified using an in-house method called cell2tissue, which is designed to expand cell masks to tissue segmentation images. The cell masks are dilated, eroded and filtered to tissue masks using skimage.morphology. The tissue masks of sections are subsequently applied to gene count matrices and cell bin data to generate tissue-cut gene count matrices and tissue-cut cell bin data, while they are available for binning in different sizes and single-cell processing procedures respectively.

### Spatial Domain Recognition at Bin50 Resolution

To identify various embryo tissues, we delineate different spatial domains in spatial transcriptomic data by binning tissue-cut gene count matrices into squares with 50 sequencing spots (DNA nanoballs) in both length and width (Bin 50 data). Counts of the same gene are aggregated within each bin using st.io.read_bgi() in Spateo^65^.

To recognize functional spatial domains, we employ data filtration, normalization, and clustering for bin50 data. To mitigate potential noise in the clustering process, only protein-coding genes expressed in ≥ 3 cells within a section and cells with total gene counts > 100 (for E9.5 embryo data) or > 200 (for E11.5 embryo data) are retained for analysis. Normalization is performed using dyn.pp.normalize_cell_expr_by_size_factors() in Dynamo^66^. Top 50 PCs are selected, and 30 neighbors are considered when computing the neighborhood graph. Unsupervised clustering is completed using st.tl.scc() in Spateo with a resolution parameter of 0.8. We visualize cluster similarity by uniform manifold approximation and projection (UMAP) in Scanpy^67^. Differentially expressed genes (DEGs) for each cluster are identified using sc.tl.ranked_genes_groups() with the Wilcoxon rank-sum test.

Using the E-mouse atlas^68,69^ as an anatomical reference for the organ and tissue distribution, we manually recognize spatial domains on clearly clustered sections by their top 30 DEGs and anatomical references. To increase the amount of data for E9.5 embryos, consecutive high-quality sections, such as S17∼S20, S37∼S40, S65∼70 and S81∼S82 are clustered and recognized in groups respectively. To reduce batch effects across above groups of sections, we also performed Harmony^70^ before unsupervised clustering.

To refine the inconclusive recognition of spatial domains on non-clearly clustered or low-quality sections, we select well identified sections as references to transfer annotations to adjacent sections using an in-house graph attention (GAT)-based method. For E9.5 mouse embryos, transferring spatial domains involves assigning S19 for S1∼S24, S37 for S25∼S62, S68 for S63∼S78, S81 for S79∼S94. While for E11.5 mouse embryos, we assign S6, S22, S35, S36, S41, S52, S57, S58, S69, S75 and S84 as references to most adjacent sections. To smooth the data, we uniformly assign bins with the dominant annotation considering their 40 nearest neighbors that have a particular annotation with a proportion greater than 80%.

All AnnData objects with annotations were saved as .h5ad files after transfer completion. Visualized bin50s with spatial domain annotations were saved as .tif images for registration and 3D reconstruction.

### Blockface Image Correction

Blockface images are designated as the gold standard for registration of sections and 3D reconstruction, as they embed the original coordinates of sections in unprocessed mouse embryos. However, tissue regions in raw blockface images are challenging to recognize.

For E9.5 mouse embryos, tissue regions in blockface images were segmented manually. For E11.5 mouse embryos, adjacent H&E images are registered to blockface images to facilitate recognition of tissue regions. The dimensions of processed images are maintained, and all pixels in non-tissue areas are removed, resulting in a refined outline of mouse embryonic tissue.

Registered ssDNA staining images are utilized to further refine the tissue regions in blockface images. For each section, the blockface image and corresponding registered ssDNA image are imported into ImageJ. The hue and saturation of registered ssDNA images are adjusted for improved distinction, and layers are merged to simplify processing. The opacity of the blockface image is adjusted to achieve optimal effect. During image registration, rotation is applied as needed to match the orientation of registered ssDNA images. Finally, blockface images are further refined by finely adjusting the brightness or darkness of specific areas through precise brushing and other operations. The modified images are saved, ensuring that dimensions meet the requirements for quality and accuracy of the final output images.

### Whole Embryo 3D Reconstruction

3D reconstruction of spatial transcriptomics data is based on the alignment of all sections from each embryo. To mitigate the impact of slight distortions caused by the slicing process, we utilize bin50 images from spatial transcriptomic data for 3D reconstruction. At first, we register each bin50 image with spatial domains to its corrected blockface image using rigid and elastic registration. Rigid registration includes flipping, translation, and rotation (scaling was not required) for rough alignments. Then elastic registration is performed to further refine the alignment by subtle distortions and deformations between images. In elastic registration, anchor points are strategically chosen from areas with clear anatomical features or boundaries in both blockface and bin50 images to ensure accurate registration. TrakEM2^71^ within Fiji^72^ is employed for these registration, while the registered images and registered transformation relationships are saved.

After registration to blockface images, we reutilize transformation relationships from registered bin50 images to cell bin coordinates. Then we merge all sections with registered coordinates to a whole 3D points cloud for E9.5 embryo and E11.5 embryo respectively. The whole 3D points cloud are input into CloudCompare (v2.13.0) to generate body mesh and remove spots outside the mouse body. With annotations of spatial domains, we also generate meshes for representative tissues and remove spots outside their anatomical structures. Then these tissue meshes are also utilized to refine annotations of spatial domains in spatial transcriptomic data by 3D points cloud. The z-coordinates are also determined in this step based on the recorded sequence of sections.

Following all 3D reconstruction procedures, x, y, z coordinates of captured cells were determined, and the mass centers of cells were saved in .h5ad files for each section.

### Digitization of Embryo along Axis

Cells from the whole embryo, brain and spinal cord are extracted based on previously refined spatial domains. To analyze the expression pattern along each axis, these cells are aligned along anterior-posterior (AP) axis and projected onto the dorsal-ventral (DV) and left-right (LR) axes.

For the whole embryo, we divide it into upper and lower body, and manually extract the dorsal and ventral median point clouds for each part. These point clouds are utilized to construct principal curves using the PrinCurve_method function in Spateo. Cells from the upper and lower body are then mapped back to their respective dorsal principal curve using KDTree (scipy) to generate new z coordinates along the AP axis. These new z coordinates are mapped onto their ventral principal curves to derive each vector A, which indicates the direction from dorsal to ventral. For each cell, another vector B is constructed from its original coordinates to its corresponding dorsal principal curve at the same z coordinates. With vector A and vector B, the radians across cells and the DV axis are calculated via cross product operations. Finally, leveraging the radians and the distance of each cell from its dorsal principal curve, all cells are projected onto DV and LR axes.

For the brain, we divide it into forebrain, midbrain and hindbrain parts, and manually extract the dorsal and ventral median point clouds for each. The same pipeline as for the whole embryo is utilized to obtain new positions of all cells along AP, DV and LR axes.

For the spinal cord, we analyze itself, and construct principal curves of the corresponding notochord. To expedite computation for E11.5 spinal cord, a down-sampled subset of 60,000 cells (out of 692,988 cells) is utilized to construct the dorsal principal curve, while all cells are utilized for E9.5 spinal cord. With the corresponding notochord, we construct the ventral principal curve for each embryo. Then the same pipeline used for the whole embryo is performed to generate new positions of all cells along AP, DV and LR axes.

### Pseudo-slices Reconstruction

To generate a pseudo-slice from any angle, we manually create a new section utilizing the point cloud of the whole embryo as input in CloudCompare. To enhance expression signals in the transverse section, specific tissue cells are projected onto a unified transverse plane across LR and DV axes following the same pipeline used in the digitization process.

### Identification of Axis Variable Genes

Three axes (AP, DV, and LR) of the whole embryo, brain, and spinal cord were identified by principal curves that pass the middle of cells smoothly. These curves are composed of discrete points that represent the projection of cells on the curve. We then split each curve into 50 sections, with each section containing the same number of points. Since each point represents the projection of a cell, all sections have the same number of cells. By averaging gene expression levels for cells within each section, we obtained gene expression patterns for all genes along any axis. Lastly, we manually selected axis variable genes and displayed them sequentially according to their highest expressed sections.

### Cell Type Annotation at Cellbin Resolution

To address the imbalance in cell counts within clusters caused by global cell annotation of the whole embryo, hierarchical clustering is employed for annotating individual cells. Unsupervised clustering is performed within each spatial domain, resulting in improved accuracy and contextually relevant annotations, thus enhancing the overall precision of the annotation process.

At first, cells within each spatial domain are extracted from the whole embryo data. To facilitate annotation by transferring cell types from single-cell datasets, spatial domains with similar cell types are merged (Table 1). The cells from each spatial domain are preprocessed using Spateo^65^ and Scanpy^67^, which involve gene and cell filtration, normalization, and unsupervised clustering. To reduce potential noise during clustering, only protein-coding genes expressed in at least 3 cells within a section and cells with total gene counts greater than 100 are retained for analysis. Normalization is performed using the recipe_pearson_residuals function in Scanpy, with 5,000 highly variable genes (HVGs) and batch correction across sections.

For E9.5 spatial domains (excluding the brain), principal component analysis (PCA) is conducted on the normalized expression matrix, followed by Harmony to further reduce batch effects across sections. Then the top 50 PCs are selected, and 30 neighbors are considered when computing the neighborhood graph. Unsupervised clustering is performed using the louvain function in Dynamo^66^ with a resolution parameter of 0.8, followed by UMAP to visualize the clustering. For the brain, scGen^73^ is employed with sub regional annotations (forebrain, mid-hindbrain, hindbrain, midbrain) to replace Harmony for batch effect removal.

For E11.5 spatial domains, only Pearson residuals are used to reduce batch effects, with other filtration, normalization, and unsupervised clustering steps following the same pipeline as for E9.5 spatial domains.

Clusters of cells within spatial domains are annotated by aligning them with a well-annotated mouse embryonic single-cell dataset^5^. The alignment is achieved by calculating Spearman correlations between aggregated expression for each gene of query and target cells to identify the best-matching cell type (Table 2 for cell types used as query in different spatial domains).

### Identification of Spatially Co-expressed Gene Modules

To identify spatially co-expressed gene modules at the microenvironment level, we divide the whole embryo into 0.05mm cubes (Bin100 data). Gene counts are aggregated within each cube, and the dominant spatial domain of each cube is assigned as its annation. Spatially co-expressed genes with each spatial domain at Bin 100 resolution are identified and clustered into modules using Hotspot.

Specifically, top 5000 highly variable genes are selected as input. Gene expression within each Bin100 is normalized by size factor. We then calculate the spatial autocorrelation score using the compute_autocorrelations function. Genes with significant spatial autocorrelation (p < 0.05) are retained and further clustered into modules using the create_modules function with a minimum gene threshold of 50 and an FDR threshold of 0.05. These identified modules are annotated to relevant anatomical structures based on their spatial localization and the co-expressed gene content.

To determine the function of each spatially co-expressed gene module, we perform functional enrichment analysis. GO enrichment score and significance levels are calculated using the clusterProfiler function with the org.Mm.eg.db database. The spatial autocorrelation scores for each module are mapped onto the whole embryo in 3D for visualization with the point cloud.

For the manually selected developmental process, we extracted the gene sets from the Mouse Genome Informatics website, and calculated geneset scores for each Bin100, using the score_genes function in Scanpy. These geneset scores for each module are similarly mapped onto the whole embryo in 3D for visualization with the point cloud.

### Identification of Ligand‒Receptor Interaction

We used Cellchat software^74^ to analyze signal communication in different tissues. Data preprocessing was performed using subsetData function, overexpressed genes were identified using identifyOverExpressedGenes, and overexpressed ligand-receptor pairs were identified using identifyOverExpressedInteractions. The overall signal communication network was obtained using computeCommunProb, and the ligand-receptor pair interaction results related to each signal pathway were calculated using computeCommunProbPathway. Visualization functions include netVisual_aggregate, netAnalysis_contribution, netAnalysis_signalingRole_heatmap, and netVisual_individual.

### Spatially resolved gene regulatory networks

SCENIC^75^ was used to analyze regulatory activity and determine the status of different tissues. The pySCENIC package in Python (version 3.8) was used for analysis to evaluate the enrichment of transcription factors and the activity of regulators. The co-expression gene network of transcription factors was calculated using the GENIE3 algorithm, using the bin100 matrix as input. The transcription factor co-expression module was identified by RcisTarget, and potential targets were filtered out using default parameters. The AUCell(area under the curve) function was used to analyze the activity of regulators, and the active regulons were determined using AUCell default parameters. The standard deviation of the regulons is calculated and visualized using the pheatmap function.

## Supplemental figures

**Figure S1.**
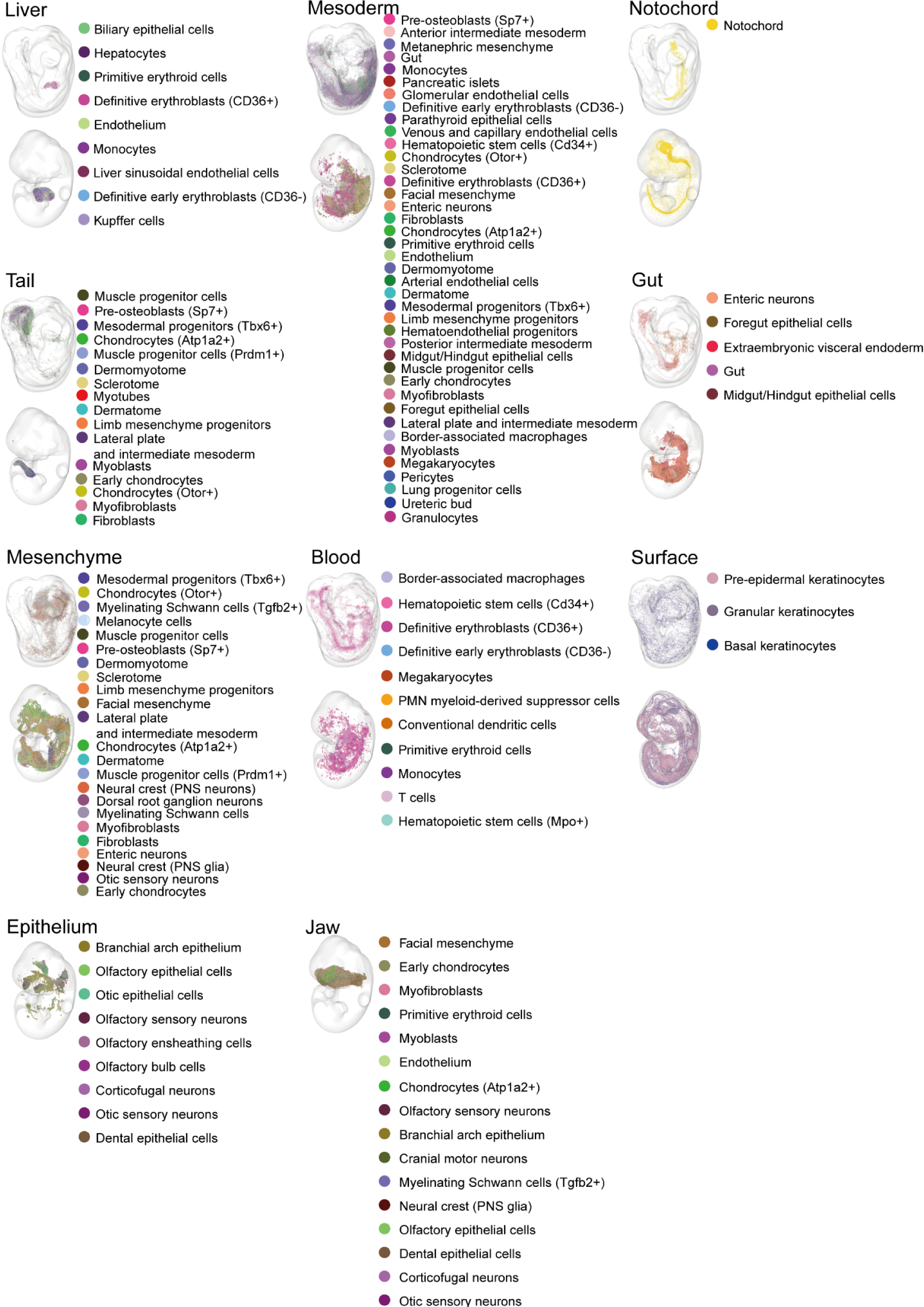
The Distribution of cell types for E9.5 and E11.5 mouse embryos, related to Figure 2. Visualization of cell type distributions for organs, including liver, mesoderm, notochord, tail, gut, mesenchyme, blood, surface, epithelium, and jaw.

**Figure S2.**
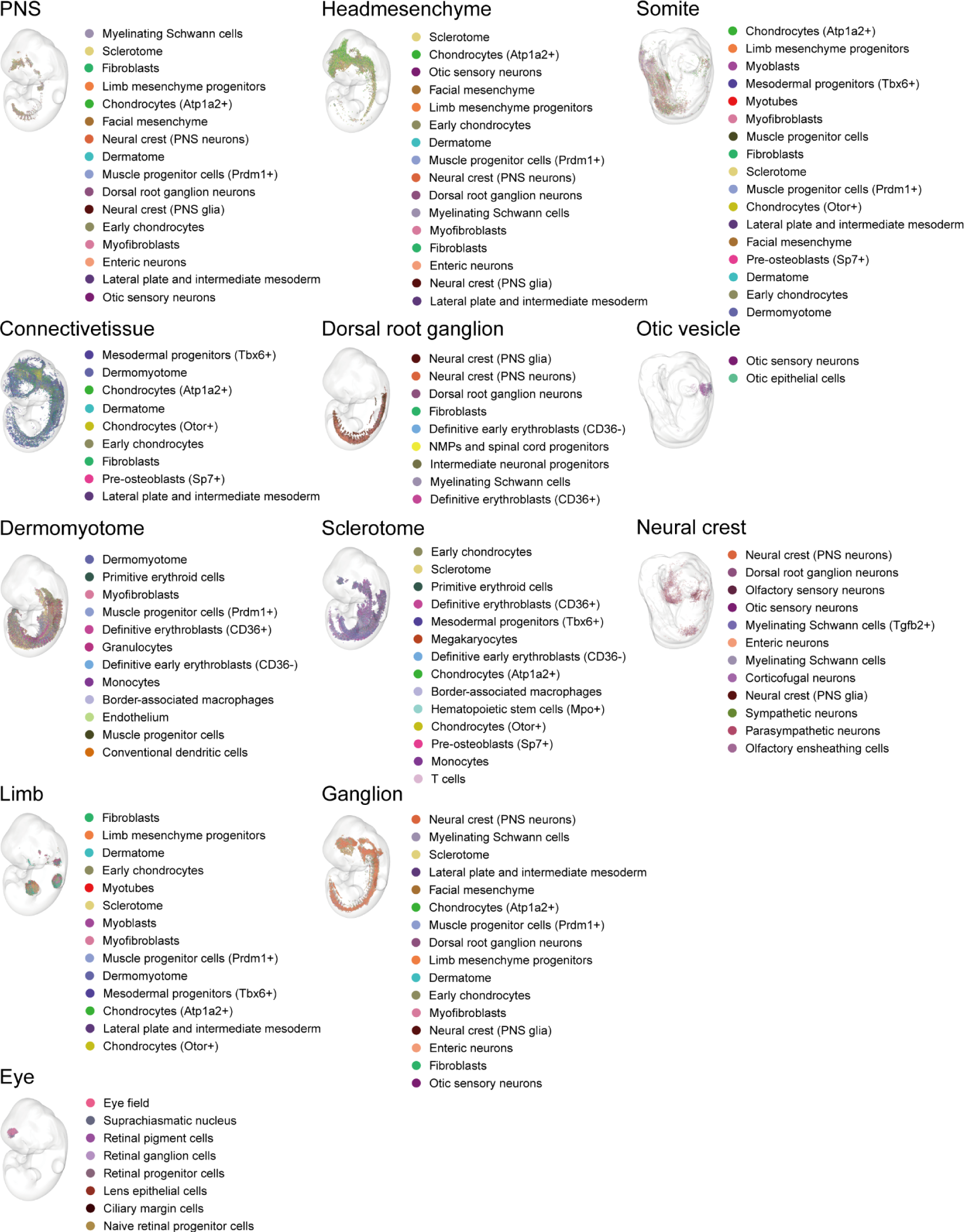
The Distribution of cell types for E9.5 and E11.5 mouse embryos, following Figure S1. Visualization of cell type distributions for organs, including sclerotome, dermomyotome, connective tissue, PNS, limb, head mesenchyme, eye, otic, dorsal root ganglion, ganglion, somite, and neural crest.

